# Effect of wall type, delayed mortality and mosquito age on the residual efficacy of a clothianidin-based indoor residual spray formulation (SumiShield™ 50WG) in southern Mozambique

**DOI:** 10.1101/2021.03.03.433723

**Authors:** Helena Marti-Soler, Mara Maquina, Mercy Opiyo, Celso Alafo, Ellie Sherrard-Smith, Arlindo Malheia, Nelson Cuamba, Charfudin Sacoor, Regina Rabinovich, Pedro Aide, Francisco Saúte, Krijn Paaijmans

## Abstract

**Background:** Indoor residual spraying (IRS) is one of the main malaria vector control strategies in Mozambique alongside the distribution of insecticide treated nets. As part of the national insecticide resistance management strategy, Mozambique introduced SumiShield™ 50WG, a third generation IRS product, in 2018. Its residual efficacy was assessed in southern Mozambique during the 2018-2019 malaria season.

**Methods:** Two different wall surfaces, cement and mud-plastered surfaces, daily mosquito mortality up to 120 hours post-exposure, and older mosquitoes (13-26d old) were included in standard WHO (World Health Organization) cone bioassay tests. Lethal times (LT) 90, LT50 and LT10 were estimated using Bayesian models.

**Results:** Mortality 24h post exposure was consistently below 80%, the current WHO threshold value for effective IRS, in both young and old mosquitoes, regardless of wall surface type. Considering delayed mortality, residual efficacies (mosquito mortality equal or greater than 80%) ranged from 1 to ≥12 months, with the duration depending on mortality time post exposure, wall type and mosquito age. Looking at mortality 72h after exposure, residual efficacy was between 6 and 9 months, depending on wall type and mosquito age. Mortality of older mosquitoes was significantly higher on mud-surfaces compared to cemented-surfaces 24h post exposure, but this difference was not significant for the delayed mortalities. The LT_50_ and LT_10_ (i.e. 90% of the mosquitoes survive exposure to the insecticides) values were consistently higher for older mosquitoes using the 24h post-exposure observations and ranged from 0.2 to 5.7 months and 0.2 to 7.2 months for LT50 and LT10, respectively.

**Conclusions:** The present study highlights the need for assessing mosquito mortality beyond the currently recommended 24h post exposure. Failure to do so may lead to underestimation of the residual efficacy of IRS products, as delayed mortality will lead to a further reduction in mosquito vector populations and potentially negatively impact disease transmission. Monitoring residual efficacy on relevant wall surfaces, including old mosquitoes that are ultimately responsible for malaria transmission, and assessing delayed mortalities are critical to provide accurate and actionable data to guide vector control programmes.

## Introduction

Indoor residual spraying (IRS) was introduced in southern Mozambique in 1946 in the semi-urban and rural parts of Maputo City and Limpopo Valley, respectively [1]. IRS with DDT (dichlorodiphenyltrichloroethane) and/or BHC (benzene hexachloride) was implemented until 1969 [1]. Following the end of the civil war in the 1990s, IRS resumed with pyrethroids (deltamethrin and lambda-cyhalothrin), which were replaced by bendiocarb in 2000 [2]. That year saw an intensification of IRS (until 2011) under the Lubombo Spatial Development Initiative (LSDI) with the aim to interrupt malaria transmission in the southern part of Mozambique [2]. While the goal to interrupt malaria was not achieved under this initiative, a significant reduction in malaria burden was observed [2]. In 2015, the MOSASWA (Mozambique, South Africa, Swaziland) initiative was launched with the aim to renew the regional efforts to accelerate progress towards malaria elimination goals already established in the region [3]. The goal was to achieve zero local transmission in Maputo Province by 2020 and to achieve pre-elimination status in Gaza and Inhambane provinces by 2025.

Mozambique’s primary vector control strategy is a countrywide universal coverage with long-lasting insecticidal nets (LLINs) [4]. IRS is used in Zambézia province [5] and southern Mozambique [6]. In both locations, non-pyrethroid insecticides are deployed to manage insecticide resistance in areas with high coverage of pyrethroid-LLINs [7]. Pyrethroid resistance in the major malaria vector *Anopheles funestus* is widespread in southern Mozambique [8–11], and recently a dramatic loss of efficacy of pyrethroid-LLINs, including those with piperonyl butoxide (PBO), was observed [12]. In addition, implementing both tools in the same geographic areas can lead to an additional reduction in malaria, although evidence is not always in agreement [13] and high costs for implementing IRS have impeded it’s use more widely.

IRS involves the application of an insecticidal product, approved for the use in public health, to the interior wall (and sometimes roof) surfaces of homes. The active ingredient kills resting mosquitoes that come into contact with these insecticide-treated surfaces. The residual efficacy of a product, or its duration of protection, is typically assessed via the World Health Organization (WHO) cone wall bioassays [14] and quantified as the number of months post-application whereby the mortality of susceptible mosquitoes is >80% [15]. This metric is important to time the start of IRS operations and/or to assess if more spray rounds during the malaria season are needed.

In 2018, Mozambique introduced SumiShield™ 50WG (50%, w/w, Sumitomo Chemical Co., Ltd., Tokyo, Japan) to its IRS portfolio, as part of its insecticide resistance management strategy. The active ingredient in this third generation IRS product is clothianidin, belonging to the chemical class neonicotinoides. It has been shown to retain its bio-efficacy for much longer compared to other IRS products [16]. However, no information exists on how this product performs on different wall surfaces in southern Mozambique. As part of routine monitoring and evaluation (M&E) operations, the residual efficacy of SumiShield™ 50WG was evaluated during the 2018-2019 malaria season. Two different -but common- wall surface (cement and mud-plastered) were evaluated, as type of surface is known to affect the residual efficacy of a variety of IRS products [17–19]. Apart from testing young (2-5 day old) mosquitoes, which is the current recommendation of WHO, older mosquitoes were exposed as well, and mortality post-exposure tracked for up to 5 days. This older mosquito cohort is important to track, as they are ultimately responsible for malaria transmission: malaria parasite development times inside the mosquito vector ranges from 9 days to several weeks, depending on temperature [20, 21].

## Materials and Methods

### Study area

The residual efficacy of SumiShield™ 50WG was evaluated in the village of Palmeira (25° 15’ 19’’S; 32° 52’ 22’’E), Manhiça district, Maputo province (southern Mozambique). Health and Demographic Surveillance System (HDSS) data collected routinely by the Manhiça Health Research Centre [22] showed the following distribution of wall types in houses in Palmeira (2019 survey): cement (11.3%), mud bricks (59.1%), cane (27.7%), and other material (1.9%). Unfortunate, these data represent the main structure of the buildings, but do not include information on the wall surface itself on which mosquitoes rest. As such the following sixteen representative houses were selected: Eight houses with mud-plastered walls (including structures with mud brick and cane walls) and eight with cement-plastered walls (including structures made with cement blocks). For each wall type, six houses were sprayed; two houses were not sprayed (owner refused spraying or was not present at the time of IRS application) and served as controls.

### IRS implementation

IRS was implemented (following the guidelines of 300 mg ai/m^2^) by the National Malaria Control Programme (NMCP) and Goodbye Malaria Initiative (Tchau Tchau Malaria) from September 1^st^ 2018 to 11^th^ February 2019 in the village of Palmeira. The entire walls of eligible structures were sprayed, as well as ceiling surfaces if these were cement-plastered or made of bricks. Residual efficacy was monitored monthly from November 2018 (month 1 post-IRS) to October 2019 (month 12).

### Residual efficacy of IRS insecticides

Cone bioassays were conducted using WHO-standard cones and exposure procedures [15]. The cones were positioned at three different heights (approx. 0.4, 1.0 and 1.6m from the ground, referred to as bottom, middle and upper, respectively) on one wall, after which ten female *Anopheles arabiensis* mosquitoes (KGB strain, details on mosquito age below) were introduced in each cone for 30 minutes. Knock-down (kd) was scored at the end of the 30-minute exposure, after which mosquitoes were transferred to paper cups and transported to the insectary facility at CISM, where they were kept in the insectary (at 25±2°C, 80±5% RH, and a12:12h day:night cycle) with *ad libitum* access to 10% sugar water (refined white sugar, sucrose). Mortality was observed 24h, 48h, 72h, and 96h and 120h post-exposure to assess (delayed) mortality.

### Susceptible mosquito line and mosquito age

*An. arabiensis* mosquitoes (CISM-KGB strain, originally obtained from Vector Control Laboratory Unit at NICD-NHLS, Witwatersrand University, South Africa) were used to assess residual efficacy. The strain is maintained at CISM’s insectary (similar conditions as described above) and its insecticide susceptibility is regularly re-confirmed. WHO tube tests [23] conducted in 2019 showed susceptibility to 4% DDT, 0.05% deltamethrin, 0.25% pirimiphos-methyl and 0.1 % bendiocarb (see S1 Table 1). Note that susceptibility to clothianidin was not tested due to a lack of standardized WHO guidelines for this particular insecticide. It is expected to show delayed mortality, killing equivalent proportions of exposed mosquitoes after 72h relative to the 24h mortality assessment with other insecticide classes [24].

Every month, 2-5 day-old unfed female mosquitoes (hereafter referred to as ‘young’ mosquitoes) were used in the wall cone bioassays. From month four onward, older and previously blood-fed mosquitoes were also tested using the same methodology. These mosquitoes were used to maintain the colony. Their age ranged from 13 to 26 days and they had been offered 3 to 5 blood meals prior to the exposure, using Hemotek membrane feeders (Discovery Workshops, Accrington, UK) with bovine blood. These females, hereafter referred to as ‘old’ mosquitoes, were allowed to oviposit and were unfed at the time of exposure. Per house, and thus wall type, bioassays with young and old mosquitoes were performed separately but during the same morning.

### Data analysis

Mortality in both insecticide exposed and control treatments was calculated as the number of mosquitoes that was dead out of the total mosquitoes tested for each of the three cones in each house. When control (exposure to no insecticide) mortality 24h post-exposure was greater than 20% (in the three cones combined), the bioassay was discarded. When control mortality was equal to or greater than 5%, mortality (for 24h and all delayed mortalities) of exposed mosquitoes was corrected using Abbott’s formula [15, 25]. Only in a single situation where the observed control mortality, for 120h delayed mortality, was 100%, mortality was not corrected as Abbott’s formula is not defined in that specific situation. To evaluate residual efficacy, the number of months during which mosquito mortality is equal to or greater than 80% is reported, following WHO guidelines [15]. Data were analyzed using R statistical software, version 3.6.1 [26]. The number of treated houses included in the analyses can be found in S1 Tables 2 and 3.

Poisson regression models were fitted to analyze differences in the observed mortality over the study period (twelve months), using the number of houses as the offset, according to cone height (lower, middle, upper), wall surface (mud, cement) and mosquitoes’ age (young, old). Here, the Abbot’s adjusted mosquito mortality (y) for each month *i* is estimated by the covariates – cone height, wall surface and mosquitoes’ age – in matrix **X**:

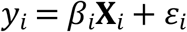

The coefficients of each term is indicated by *β,* the error term is indicated by *ε* which accounts for variability in the data that is not explained by the covariates. Models were fitted using maximum likelihood (mixlm package [27])

Logistic binomial models were fitted through actual residual bio-efficacy data, such that:

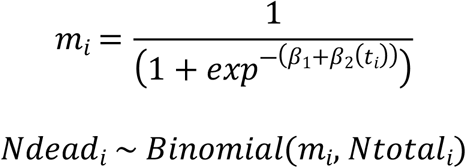

where *m* indicates the probability that a mosquito is killed, *Ndead* indicates the absolute number of mosquitoes killed per sample *i* and *Ntotal* is the number tested which is assumed to have a logistic distribution over time *t.* The parameters *β*_*1*_ and *β*_*2*_ determine the duration and depreciation of the logistic fit. Bayesian models were fitted using Hamiltonian Monte Carlo sampling methods [51, 52]. Four chains were initialized to assess the convergence of 1000 iterations, the first 500 of each were discarded as burn in. The posterior distributions of parameters (4000 iterations) and 90% Bayesian credible intervals were estimated, posterior checks were performed using shinystan (version 2.50.0) [28, 29] and visually confirmed to fit the data. The model was used to estimate lethal times for 90%, 50% and 10% mortalities (LT90, LT50 and LT10, respectively) for all mortality time points, and each wall type.

### Ethical considerations

Ethical approval was obtained from Manhiça Health Research Centre Institutional Bioethics Committee for Health (CIBS-CISM/006/2018; CIBS-CISM/065/2019. The household owner (>18 years old) where the WHO cone bioassays were performed was informed about the purpose of the study in the local language (Shangana or Portuguese) and gave their oral informed consent. They were free to withdraw from the study at any time.

## Results

### Effect of cone height and house replicate on mosquito mortality

There was no effect of cone height on 24h mortality on cement-plastered walls for young: bottom (711 mosquitos, 71 house visits) vs. middle (709 mosquitoes, 71 house visits), *P*=0.862); bottom vs. upper (708 mosquitoes, 71 house visits), *P*=0.931; and middle vs. upper (*P*=0.605); and old mosquitoes: bottom (479 mosquitos, 48 house visits) vs. middle (479 mosquitoes, 48 house visits), *P*=0.910; bottom vs. upper (480 mosquitoes, 48 house visits), *P*=0.966; and middle vs. upper, *P*=0.944. The same was observed on mud-plastered walls for young: bottom (692 mosquitos, 69 house visits) vs. middle (691 mosquitoes, 69 house visits), *P*=0.986; bottom vs. upper (691 mosquitoes, 69 house visits), *P*=0.971; and middle vs. upper, *P*=0.986; and old mosquitoes: bottom (484 mosquitos, 48 house visits) vs. middle (479 mosquitoes, 48 house visits), *P*=0.948; bottom vs. upper (482 mosquitoes, 48 house visits), *P*=0.942; and middle vs. upper, *P*=0.994. As such, data from the different cone heights are grouped per house, month and mosquito age group.

### Observed residual efficacy of SumiShield™ 50WG by wall type and mosquito age

Regarding young mosquitoes, 24h mortality post-exposure (PE) was never above WHO’s 80% threshold (Fig 1, S1 Table 2), which is understandable for this product that is expected to have a delayed mortality-inducing effect with mortality rates above 80% expected at 72h PE [24]. Considering delayed mortality, the time during which mortality was equal to or greater than 80% was 1 month for 48h PE (but note it was 96% in month 5), 6 months for 72h mortality PE (but 76% in month 2), and 9 months for 96h (with 77% in month 8) and 120h mortality PE on the cement wall surfaces. On mud surfaces these values were 0 months for 24h mortality PE, 1 month for 48h mortality PE, 7 months for 72h mortality PE (but 49% in month 4 and 75% in month 5), 9 months for 96h mortality (with 55%, 79% an 78% in months 4, 5 and 8, respectively) and 120h mortality PE (with 67% in month 4). Both 24h mortality and delayed mortalities did not differ for mud-plastered and cement-plastered surfaces throughout the study period (mud: 1624 mosquitos, 54 house visits, concrete: 1620 mosquitoes, 54 house visits): *P=*0.774, *P=*0.899, *P=*0.962, *P=*0.893 and *P=*0.977 for 24h, 48h, 72h, 96h and 120h mortality, respectively).

**Fig 1.**
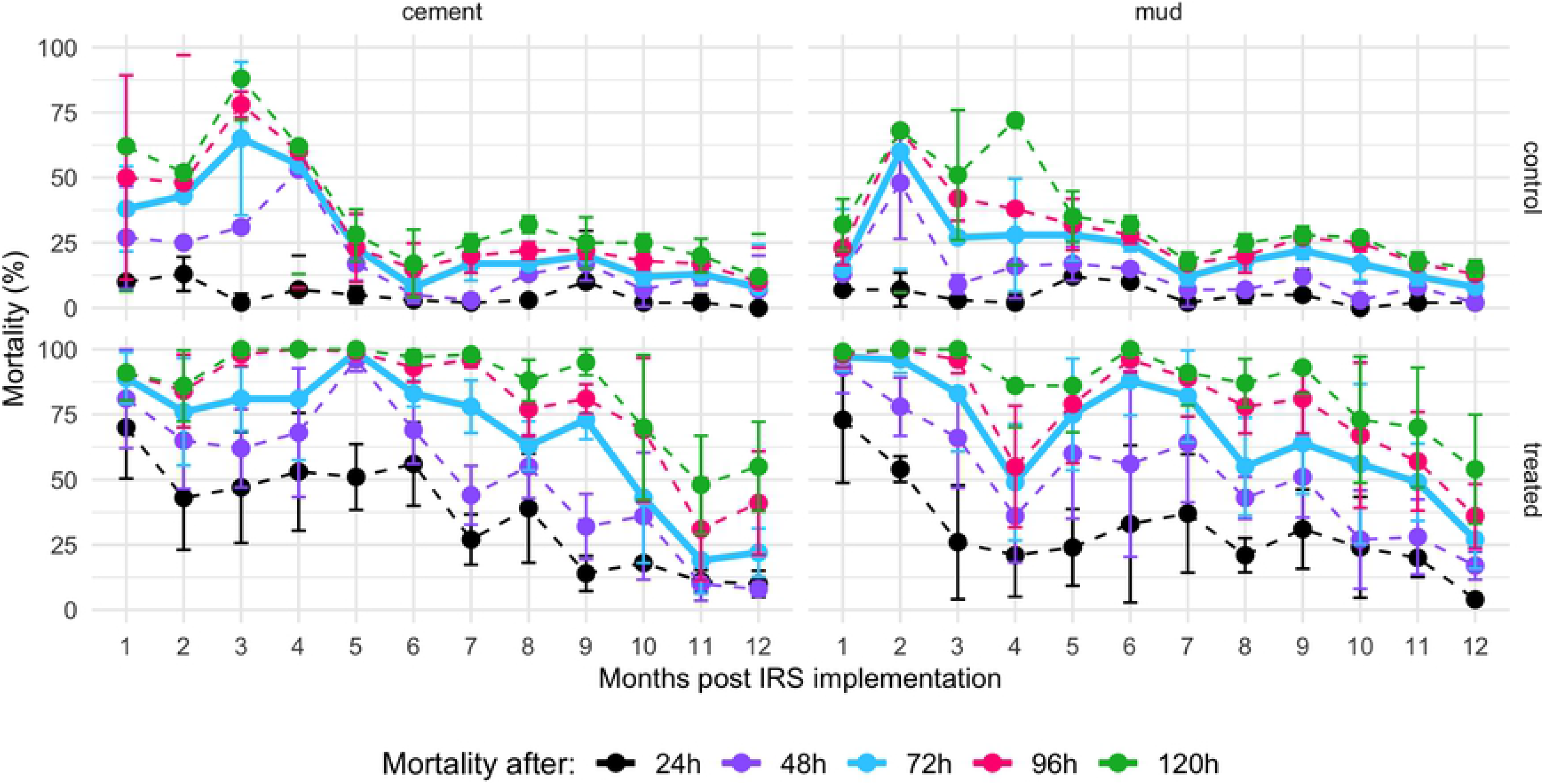
Residual efficacy of SumiShield™ 50WG on cement (left panels) and mud-plastered walls (right panels) in southern Mozambique using young (2-5d old) susceptible *An. arabiensis* female mosquitoes. Mortality was recorded 24h, 48h, 72h (highlighted as bolder blue line), 96h and 120h post a 30-min exposure to control (top panels) or treated walls (bottom panels). Mortality 72h after exposure to SumiShield™ 50WG is above 80% for 6 and 7 months on concrete and mud surfaces, respectively.

Looking at the older mosquitoes, 24h mortality was again, as expected, never above WHO’s 80% threshold (Fig 2, S1 Table 3). Considering delayed mortality, the time during which mortality was equal to or greater than 80% was 5 months for 48h, 6 months for 72h PE, and 9 months for 96h PE and 120h PE on the cement wall surfaces. On mud surfaces these values were 0 months for 24h and 48h PE, 9 months for 72h mortality PE (with 62% in month 8), 10 months for 96h mortality PE and ≥12 months for 120h mortality PE. Both 24h mortality and delayed mortalities did not differ for mud-plastered and cement-plastered surfaces during the study period (mud: 1438 mosquitos, 48 house visits; cement: 1445 mosquitoes, 48 house visits): *P=*0.787, *P=*0.923, *P=*0.971, *P=*0.899 and *P=*0.968 for 24h, 48h, 72h, 96h and 120h mortality, respectively.

**Fig 2.**
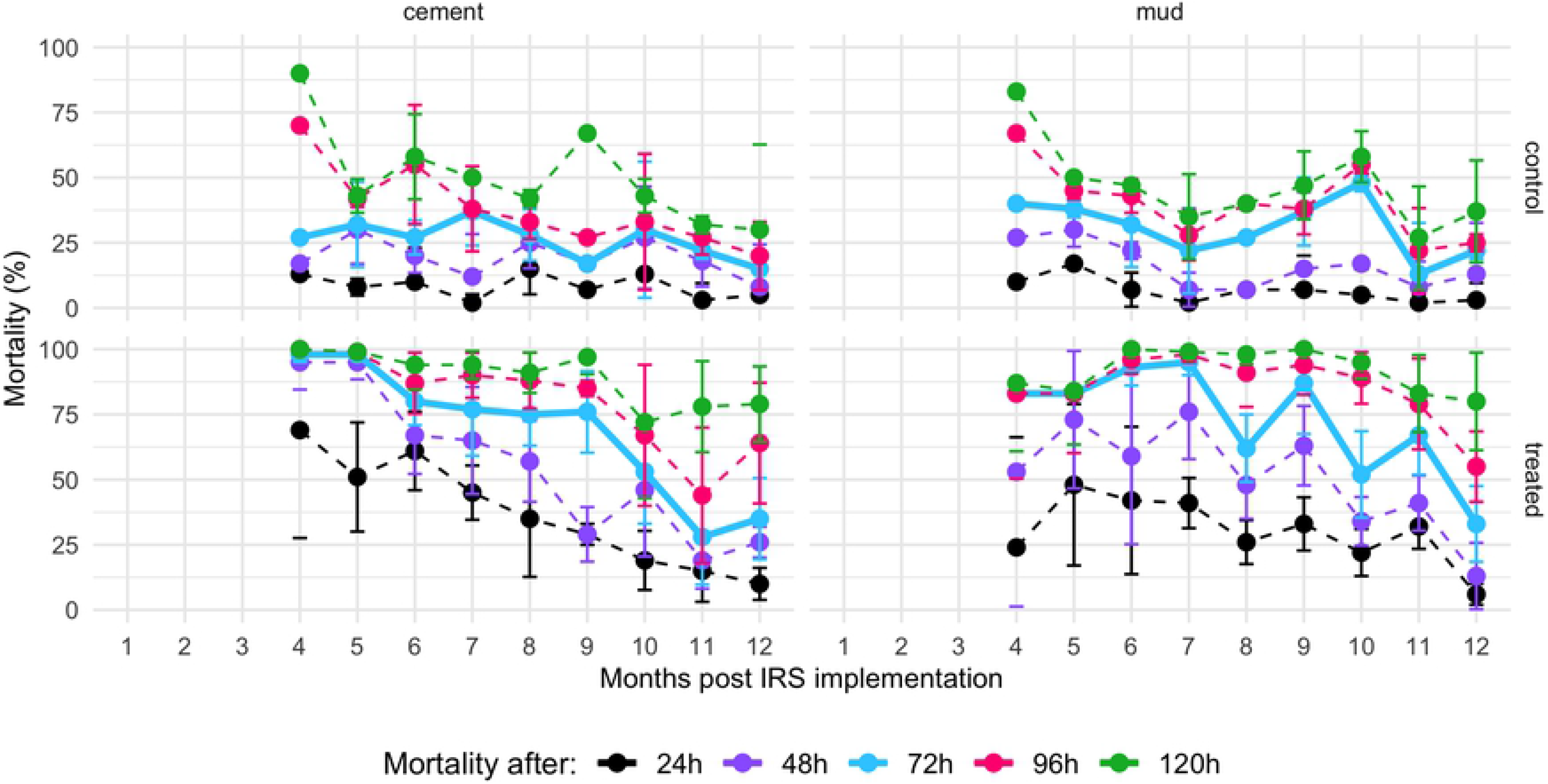
Residual efficacy of SumiShield™ 50WG on concrete-(left panels) and mud-plastered walls (right panels) in southern Mozambique using old (13-26d old) susceptible *An. arabiensis* female mosquitoes. Mortality was recorded 24h, 48h, 72h (highlighted as bolder blue line), 96h and 120h post a 30-min exposure to control (top panels) or treated walls (bottom panels). Mortality 72h after exposure to SumiShield™ 50WG is above 80% for 6 and 9 months on concrete and mud surfaces, respectively.

Comparing the mortality distribution of young and old mosquitoes on both wall surfaces, 24h mortality and delayed mortalities did not differ between young and old mosquitoes on mud-plastered (young: 1624 mosquitos, 54 house visits, old: 1445 mosquitoes, 48 house visits): *P=*0.492, *P=*0.396, *P=*0.296, *P=*0.238 and *P=*0.257 for 24h, 48h, 72h, 96h and 120h mortality, respectively) and cement surfaces (young: 1620 mosquitos, 54 house visits, old: 1438 mosquitoes, 48 house visits), *P=* 0.361, *P=*0.304, *P=*0.247, *P=*0.276 and *P=*0.252 for 24h, 48h, 72h, 96h and 120h mortality, respectively).

### Estimated residual efficacy of SumiShield™ 50WG by wall type and mosquito age

Median lethal times (LT50s) predicted by Bayesian models ranged from 10.0 to 13.1, 11.3 to 15.8 and 12.2 months to 19.3 for 72h, 96h and 120h PE, respectively, with differences depending on mosquito age and wall substrate (Fig 3 and SI Table 4). Estimated LT10 values (i.e. when residual efficacy is 10%, or when 90% of the mosquitoes survive exposure to the insecticides) ranged from 15.1 to 19.6, 15.1 to 22.3 and 15.4 to 26.3 months for 72h, 96h and 120h PE, respectively. Both LT50 and LT10 values were consistently higher for older mosquitoes, with differences ranging from 0.2 to 5.7 months and 0.2 to 7.2 months for LT50 and LT10, respectively, depending on the delayed mortality time. The mean 72h mortality in the control treatments across the 12 sampling months was 42.8% (SD: 22.3%) and 41.8% (SD: 22.3%) lower than the mortality in the exposed mosquitoes on cement-plastered and mud-plastered walls, respectively. This indicates that the higher mortality of older mosquitoes is partly due to the action of the IRS exposure, and not due to mosquito senescence alone.

**Fig 3.**
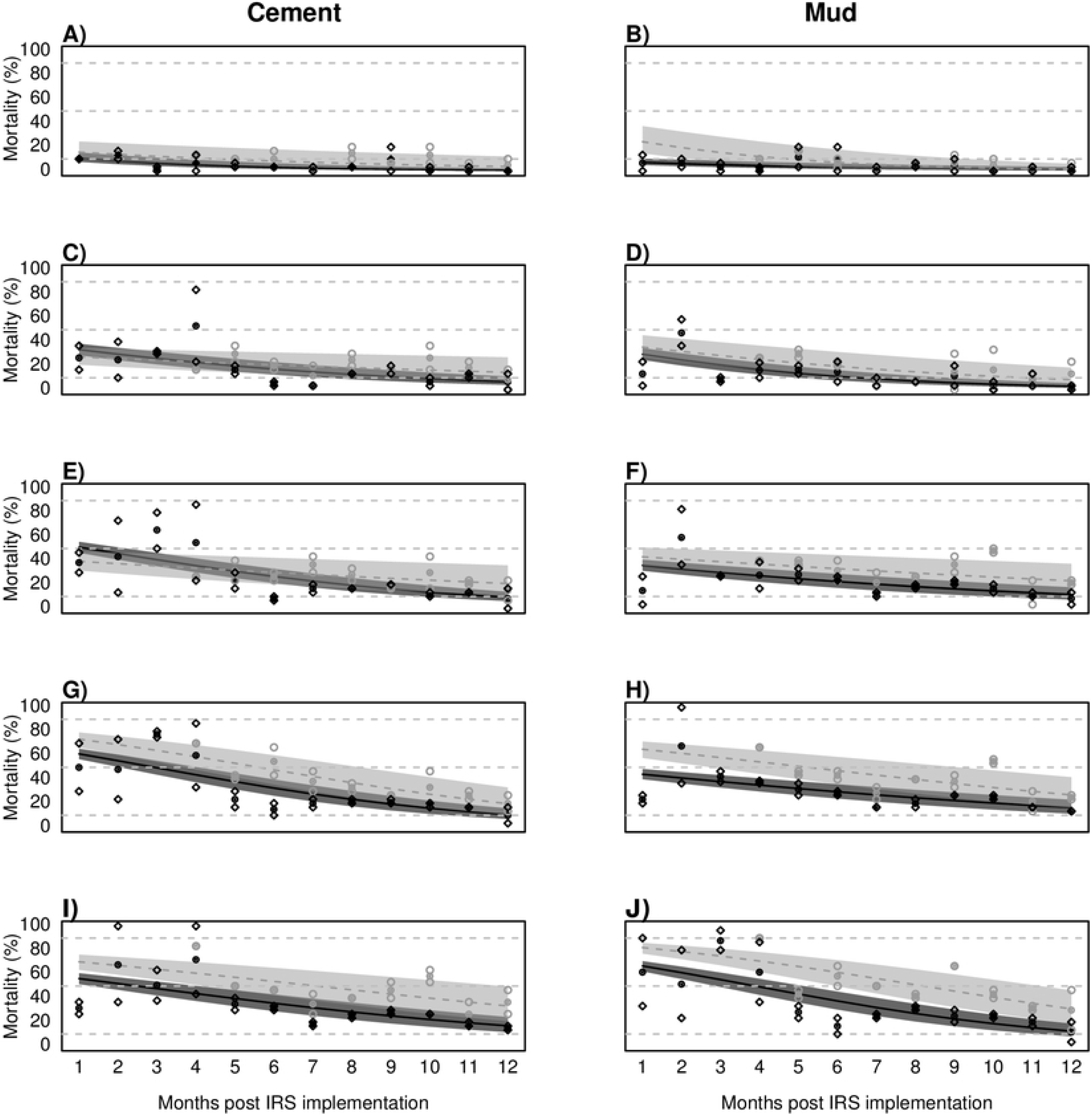
Observed and predicted delayed mosquito mortality of young (2-5d old) and older (13-26d) mosquitoes after exposure to SumiShield™ 50WG on cement (panels on the left) and mud-plastered walls (panels on the right). Data points represent observed mortality for each cone bioassay conducted, combining cone heights (6 houses); young mosquitoes are represented in dark grey; old mosquitoes in light grey. **A,B)** 24h mortality. **C,D)** 48h mortality. **E,F)** 72h mortality. **G,H)** 96h mortality. **I,J)** 120h mortality. Horizontal dashed lines in each panel indicate the 90%, 50% and 10% mortalities that were used for the lethal time estimates.

## Discussion

The 24h mortality of young (2-5 days) and old (13-26 days) laboratory-reared mosquitoes that were exposed to walls sprayed with SumiShield™ 50WG, was consistently below the WHO threshold of 80% [15] for both cement and mud-plastered surfaces. These findings corroborate previous studies that have documented the delayed killing effect of SumiShield™ 50WG [16, 30-32] and confirm that its active ingredient, clothianidin, is slow acting. Clothianidin is a neonicotinoid and acts on the insect nicotinic acetylcholine receptor [33], which in turn causes systemic toxicity upon contact and poison the mosquito gut [34, 35]. It is believed that this mechanism may be responsible for extended delayed mortality effect, which were also observed in the present study.

Considering delayed mortality (in this study daily mortality up to five days post exposure), residual efficacies (with a mosquito mortality equal or greater than 80%) ranged from 1 to ≥12 months, with the duration depending on mortality time post exposure, wall type and mosquito age. Looking at mortality 72h after exposure, residual efficacy was between 6 and 9 months, depending on wall type and mosquito age. This study confirms that failure to assess this delayed mortality (i.e. following WHO’s current guidelines and assess 24h mortality post-exposure only [14]) will lead to an underestimation of the efficacy of SumiShield™ 50WG. Furthermore, this additional (delayed) mortality will lead to a further reduction in mosquito vector populations and potentially disease transmission risk, compared to considering 24h mosquito mortality alone [30–32].

We show here that mosquito age can affect the efficacy of vector control products with mosquito mortality being higher in older mosquitoes [36, 37]. The extent to which mosquito age plays a role likely depends on the type of active ingredient used. The present study demonstrated that SumiShield™ 50WG kills older malaria mosquitoes effectively. The 72h mortality in the control mosquitoes that were not exposed to an insecticide was on average 42-43% lower than the mortality in the exposed mosquitoes. This indicates that the higher mortality of older mosquitoes is partly induced by exposure to SumiShield™ 50WG, and not due to mosquito senescence alone.

Having said that, the age range of the older mosquitoes varied from 13 to 26 days. In addition, those mosquitoes had the opportunity to blood feed twice per week (3 to 5 times before the bioassays), but it was not recorded if and how many times those females actually took a blood meal. These are clearly areas for improvement in future studies, but we argue that a proper understanding of the impact of IRS products on older mosquito cohorts is important, as this cohort is ultimately responsible for malaria transmission. The extrinsic incubation period (EIP; the time between a mosquito becoming infected with malaria and her becoming infectious) ranges from 9 days to several weeks, depending on temperature [20, 21]. In addition, there are extra delays, as it takes one or more days between eclosure (from pupa) to a female’s first bloodmeal [38], and a mosquito may not become infected during her first blood meal. All these factors combine to determine at what age a mosquito actually becomes a malaria vector.

IRS in itself already affects the age structure of mosquito populations. There is strong evidence that the parity rate, which is the proportion of parous mosquitoes (those that have taken a blood meal and have laid eggs at least once) out of the total number of mosquitoes, declines after IRS implementation, meaning younger mosquitoes are more abundant [39–41]. Changes in the mosquito age structure, combined with a higher killing efficiency of older mosquitoes, will drastically change malaria transmission risk as the vectorial capacity is reduced [42].

Residual efficacy differed slightly between type of wall surface. This has been shown for SumiShield™ 50WG and other IRS products before, but the direction of effect is not always consistent [17–19]. A multi-country study conducted by Oxborough and colleagues across west and east Africa assessing the residual effect of different IRS products on different wall surfaces showed that residual efficacy of each IRS product was affected by the type of surface and that variations in retention of the active ingredient can differ from country to country, even on similar surfaces [43].

IRS has contributed significantly to the gains in malaria control [44] and will likely continue to play a critical role in malaria control and elimination strategies in Mozambique, especially since IRS with different non-pyrethroid insecticides can manage insecticide resistance in areas with high ITN coverage [7]. Pyrethroid resistance is widespread in *Anopheles funestus* mosquitoes, the major malaria vector, in southern Mozambique [8–11], and recently a dramatic loss of efficacy of all long-lasting insecticidal nets (LLINs), including piperonyl butoxide (PBO)-based nets was observed [12]. However, the success of IRS programs depends on a variety of factors, and not on the residual efficacy of the products alone. Those factors include the effects of different wall substrates on residual efficacy [this paper, 19, 45, 46], the quality of the product and/or its spray application [47], community acceptability before/during IRS implementation [48, 49], wall modifications after IRS implementation [50], mosquito susceptibility to the active ingredients used [51, 52], and (changes in) vector behaviors as IRS targets indoor resting mosquitoes [53, 54]. As such, several Monitoring and Evaluation activities are needed before, during and after IRS implementation.

Having said that, SumiShield™ 50WG is a welcome product in Mozambicans vector control tool kit, as its residual efficacy is longer than that of other products [16, 18, 55, 56], and it is a product that can be used in rotation with products containing a different active ingredient to slow down the emergence and spread of insecticide resistance. Adequate insecticide resistance management remains critical, as resistance to most approved IRS insecticides has been documented in sub-Saharan Africa [51, 52, 57] and there are only five approved chemical classes of IRS insecticides. Worryingly, resistance to SumiShield™ 50WG’s active ingredient, clothianidin, has been detected in Cameroon [58], which means that proper resistance monitoring, cross-border insecticide resistance management initiatives, and alternative non-insecticidal tools [59] are urgently needed in Mozambique.

## Acknowledgments

We acknowledge the support from the Spanish Ministry of Science and Innovation through the “Centro de Excelencia Severo Ochoa 2019-2023” Program (CEX2018-000806-S), and support from the Generalitat de Catalunya through the CERCA Program. CISM is supported by the Government of Mozambique and the Spanish Agency for International Development (AECID). We thank members of the Palmeira community for allowing us to monitor residual efficacy of IRS products in their homes, and GoodBye malaria for working closely with us during household selection. We also thank all the CISM entomology members for their assistance during the study.

## Supporting information captions

**S1 Table 1. Insecticide susceptibility of the *An. arabiensis* KGB colony maintained at the Manhiça Health Research Centre.** Percentage indicates percent mortality 24h following 1h exposure in the WHO tube assay; number between parentheses indicates the number of mosquitoes tested.

**SI Table 2. Percent monthly mosquito mortality 24 to 120h post-exposure to SumiShield^®^ 50WG on cement and mud-plastered walls in southern Mozambique, using young (2-5d old) susceptible** *An. arabiensis* **females**. A total of 30 mosquitoes are exposed per house, 10 mosquitoes at each height on the wall, for 30 minutes. The results shown are aggregated across these bioassays and adjusted for control mortality (10 mosquitoes at each wall height tested on unsprayed surfaces).

**SI Table 3. Percent monthly mosquito mortality 24 to 120h post-exposure to SumiShield^®^ 50WG on cement and mud -plastered walls in southern Mozambique, using older (13-26d old) susceptible *An. arabiensis* females**. A total of 30 mosquitoes are exposed per house, 10 mosquitoes at each height on the wall, for 30 minutes. The results shown are aggregated across these bioassays and adjusted for control mortality (10 mosquitoes at each wall height tested on unsprayed surfaces).

**SI Table 4. Estimated lethal times for 90%, 50% and 10% mortalities (LT90, LT50 and LT10, respectively) of young (2-5d old) and older (13-26d) mosquitoes 24 to 120h post-exposure to SumiShield^®^ 50WG on cement and mud -plastered walls in southern Mozambique**. Lethal times are given in months post-exposure.

